# BinBencher: Fast, flexible and meaningful benchmarking suite for metagenomic binning

**DOI:** 10.1101/2024.05.06.592671

**Authors:** Jakob Nybo Nissen, Pau Piera Lindéz, Simon Rasmussen

## Abstract

New methods for metagenomic binning are typically evaluated using benchmarking software, and become tuned to maximize whatever criterion is measured by the benchmark. Subtleties in benchmarking procedures can cause misleading evaluations, derailing method development. Differences between procedures used to evaluate binning tools make them hard to compare, which slows progress in the field. We introduce BinBencher, a free software suite for benchmarking, and show how BinBencher produces evaluations that are more biologically meaningful than alternative benchmarking approaches.

## 1. Introduction

In the last decade, the number of known microbial species have exploded, largely due to culture-independent methods of discovery, where genomes are reconstructed from nucleotide sequences obtained directly from environmental samples, as done in e.g. [1] and [2]. In a typical workflow, reads from an environmental sample are *de novo* assembled to contigs. Even when using the latest metagenomic assemblers with long, accurate reads, genomes are often incompletely assembled, and may be fragmented in several contigs [3], [4] To reconstruct the original genomes, the contigs can be grouped by their genome of origin, in a process called ‘binning’.

Programs used for binning, or ‘binners’, typically rely on sequence features which correlate probabilistically between sequences of the same genome, such as k-mer composition and co-abundance. Therefore, binning is an error-prone process, where most output bins usually does not accurately correspond to genomes, but have some degree of incompleteness or contamination. In attempts to improve binning accuracy, there has been a lot of papers published the last years presenting new techniques. We know of at least 19 binners published the last decade, not including updates to existing binners [5]–[23], [3].

Typically, the accuracy of a binner is measured by running the binner on a dataset with a known ground truth reference, usually an *in silico* simulated metagenome or an artificially produced mock community, and comparing the output bins against the reference, as done in e.g. [8], [9], [16]–[19], [22], [23]. An alternative approach is to directly evaluate the bins, in the absence of a ground truth reference, using statistical models. Most commonly used is CheckM[24] and its newer version CheckM2[25]. These programs cannot fully replace simulated microbiomes for several reasons: First, they are themselves calibrated using simulated data. Second, as statistical models, they are not entirely accurate and may mispredict the accuracy of a bin. Third, in the case of CheckM2, its use of machine learning makes its results less explainable than ground-truth based benchmarking, making it less useful as a guide to develop new binning techniques.

In most recent papers presenting new binners, the authors claim superior binning accuracy over their competitors in benchmarks - claims, which often conflict with those made in other papers. For example, the authors of MetaBAT showed it was more accurate than MaxBin[8], whereas [9] showed that MetaBAT was better on one dataset, but worse on another. In [16] and [10], however, MaxBin beat MetaBAT. Comparing the respective upgrades, MetaBAT2 to MaxBin2, the former was the better choice according to [17]–[19], but they perform about equally well in [22], and MaxBin2 won in [21]. Similarly, MetaBAT2 was *much* better than VAMB according to [18], and somewhat better in [21], but only on par in [22]. In contrast, [19] found VAMB to beat MetaBAT2 and, unsurprisingly, so did we in the original VAMB paper [17].

This status quo is bad for users, who can’t easily tell which binners really are the best to use, and who rarely have the time to undergo a detailed, systematic study of the many available binners. It’s also bad for tool developers, because the conflicting claims of accuracy makes it difficult to know which techniques are promising to develop further, and even to know whether the field is making progress in the sense that binners are getting more accurate over time.

Binning is not unique in being a well-studied computational problem with a proliferation of candidate evaluation techniques - parallels can be drawn to the protein folding problem, and the problem of computer vision. Both fields have benefitted greatly from standardized evaluations provided by e.g. CASP[26] and ImageNet[27], respectively. In the field of binning, the Critical Assessment of Metagenome Interpretation (CAMI) [28] and CAMI2[29] is a similar initiative that aims to standardize benchmarking of various metagenomic tools, including binners. To this end, they have developed the binning benchmarking tool AMBER[30].

In this paper, we will demonstrate how subtle differences in the benchmarking procedure have profound impact on the assessment of bins. We show how seemlingly straightforward procedures, including that used by AMBER, can result in misleading scores. We present Bin-Bencher, a benchmarking suite for binnings of simulated metagenomes which produce more biologically meaningful results.

## 2. Results

### 2.1. BinBencher can selectively include or disregard microdiversity

Creators of synthetic datasets may include distinct genomes that are highly similar in order to test how binners handle microdiversity. For example, the GI dataset (see Methods) contains sequences from 98 species that have more than one genome in the dataset. Of these, 33 species have a mean pairwise average nucleotide identity (ANI) between its genomes above 99%, and 12 above 99.9%, according to FastANI v1.34 [31] (Figure 1). In total, there are 3,371 genome pairs with an ANI ⋝ 99%, and 1,903 pairs ⋝ 99.9%.

**Figure 1:**
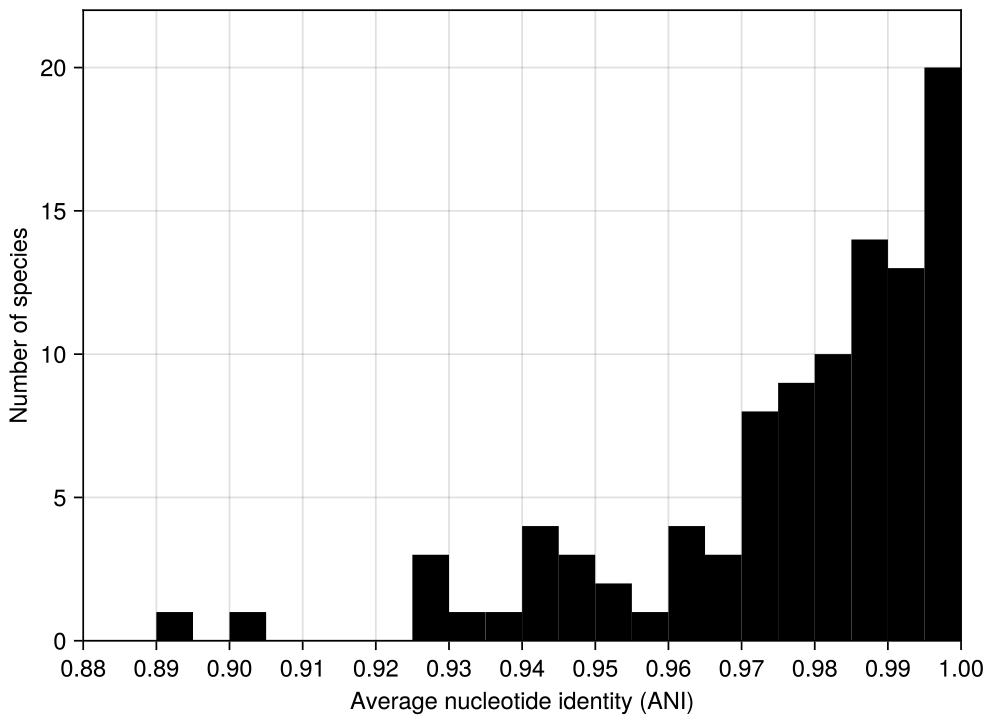
Mean average nucleotide identity (ANI) within each multi-genome species in the GI dataset. For every species with more than one genome, the mean ANI was computed across all genome-genome pairs in that species. The high level of ANI implies that computing precision at genome level can result in intra-bin microdiversity being classified as contamination despite being > 99% identical on the nucleotide level.

In this paper, we operationally define ‘microdiversity’ to be a collection of sequences from different genomes with an ANI of ⋝ 99%. Depending on the objective of the researcher performing the benchmark, microdiversity may not be considered contamination at all, but rather natural variation within a single genome. CheckM, for example, reports the fraction of contamination that results from sequences with an amino acid identity ⋝ 90% as ‘strain heterogeneity’[24]. Given that short-read assemblers struggle with preserving diversity at a >96% level [32], and the popular assembler metaSPAdes even explicitly aims to assemble the consensus of mixed strains [33] and thus co-assembles similar strains, we found it unlikely that researchers with short-read data will have microdiversity correctly preserved in the contigs input to binning.

When computing bin precision relative to a genome, sequences from any other genome count as equally contaminating, no matter if they are from a remotely related organism, or are microdiversity. If the researcher does not want to distinguish microdiversity, the bin precision can therefore be significantly underestimated. However, if the research objective is to correctly separate micro-diversity to resolved strains, then microdiversity in a bin ought to be considered contamination.

BinBencher addresses these conflicting requirements twofold: First, BinBencher allows contigs to map to any number of underlying genomes, and therefore works correctly when given bins containing contigs from a consensus assembly of different genomes. Second, BinBencher benchmarks on multiple taxonomic ranks simultaneously (see Methods). Thus, the user may define a taxonomic rank that groups highly similar genomes in their dataset, and when benchmarking, the user can choose between BinBencher’s metrics at the genome level, or at this higher taxonomic rank.

To illustrate this, of the 109 bins of the GI dataset with recall ⋝ 0.9 and nonzero contamination, 31 had zero contamination when disregarding microdiversity. However, when benchmarking on the taxonomic level of species, none of the reported contamination from any of the 68 contaminated bins with recall ⋝ 0.9 was due to microdiversity (Figure 2), because genomes from different bacterial species usually differ by more than 5% ANI[31].

**Figure 2:**
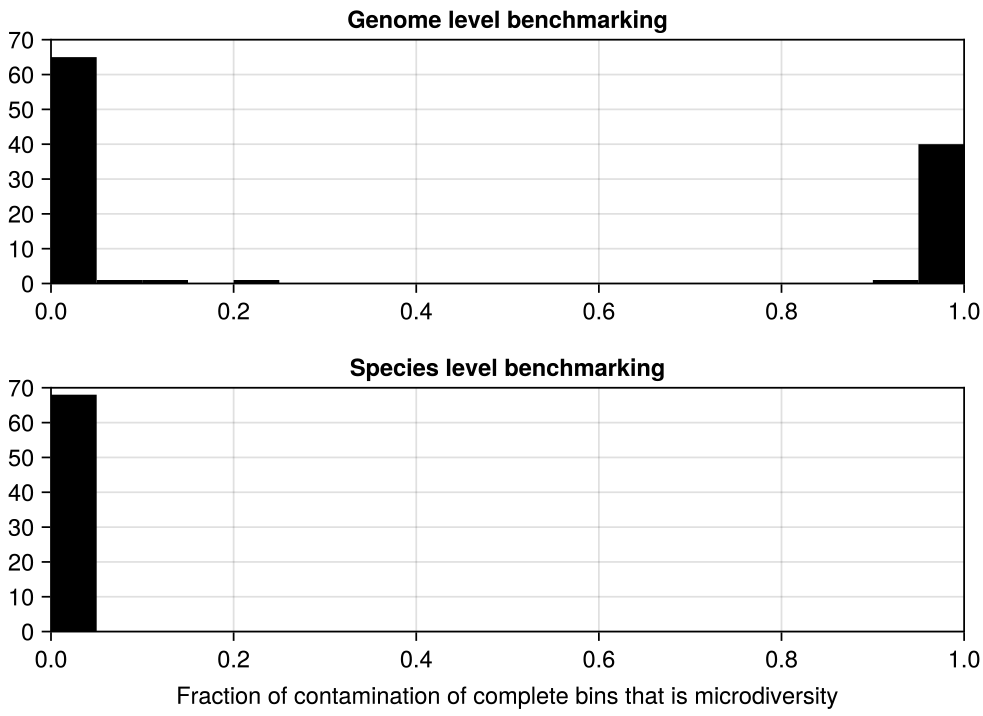
The fraction of contamination that is due to microdiversity, i.e. contamination from genomes ⋝ 0.99 ANI to the main genome in the bin. Only bins with recall ⋝ 0.9 and nonzero contamination are included. Top: Recall/precision is computed on the genome level, bottom: on the species level. The bottom plot contains less data because fewer bins have nonzero contamination on the species level. By choosing the taxonomic rank of benchmarking, users can selective choose to include or disregard microdiversity.

### 2.2. BinBencher avoids common pitfalls when evaluating multi-sample binnings

When benchmarking binnings, the same genome may be present in multiple samples. In the common ‘multi-split’ binning workflow, reads from each sample are assembled independently, contigs from all samples are binned together, and then the resulting bins are split by their sample to sample-wise pure bins. This workflow allows co-abundance to be leveraged across samples more effectively by the binner, while also resulting in bins that, because they originate from only a single sample, do not mix different strains occurring in different samples. It has been shown previously[17], [34] that multi-splitting produces more accurate bins than binning samples independently and pooling the result.

When benchmarking multi-sample binnings, the user may choose to benchmark once per sample against a sample-specific reference, or once against a single cross-sample reference. Further, the user may benchmark the output of the binner including sequences from all samples, or subset the bins by sample. These two dilemmas leads to a total of four options when benchmarking. As we will show, three of these four possibilities introduce bias when using benchmarking frameworks that compute accuracy based on the number of basepairs present in the bins relative to the reference, such as AMBER.

#### 2.2.1. Cross-sample references cause missing sequences to count against recall, even if they are redundant

If the user benchmarks against a cross-sample reference, then recall will be underestimated by AMBER. Suppose a 3 Mbp genome is present in five samples. To reach a recall of 1, a bin must have all 3 ∗ 5 = 15 Mbp present. A bin with a full 3 Mbp copy of the genome will have a reported recall of 0.2, despite the missing 12 Mbp being fully redundant, as it maps to the same genomic positions as the extant 3 Mbp.

In the GI dataset, of the 172 bins ⋝ 200 kbp with AMBER recall < 1, for 101 of them, more than half of the genomic content reported as missing by AMBER was redundant. 34 of them fully recovered the genome, i.e. all missing content was redundant (Figure 3, top). This problem was exacerbated in the multi-split workflow, because no output bins contained sequences from multiple samples, and so for 253 of the 294 incomplete bins, more than half their missing content was redundant, and for 52 of them, all missing content was redundant. (Figure 3, bottom).

**Figure 3:**
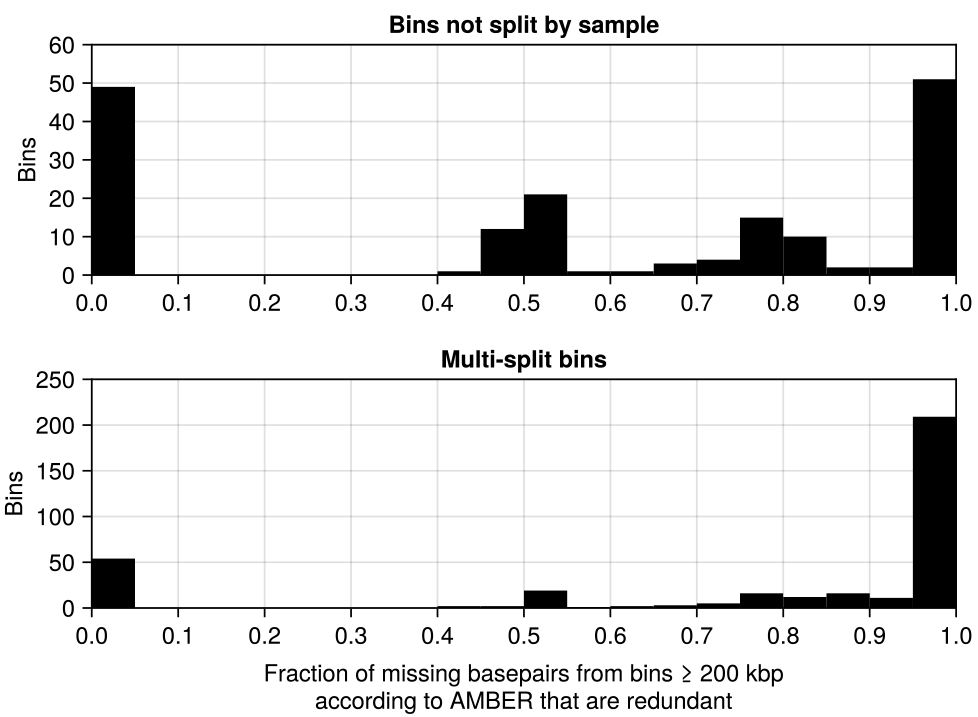
A bin’s missing basepairs (bp) is the number of bp needed for that bin to reach a recall=1 according to AMBER. We consider missing bp to be redundant if they map to the same genomic positions as bp already in the bin. Top: Most bins ⋝ 200 kbp are reported to be incomplete by AMBER, but most of the missing bp in these incomplete bins are redundant. Bottom: This effect is even more pronounced for bins from the multi-split binning workflow.

Because BinBencher computes recall in terms of genomic positions, redundant sequences are never included in the recall computation, and this issue does not occur.

#### 2.2.2. Sample-specific references cause miscomputed precision and recall for cross-sample binnings

If the user benchmarks against sample-specific references using AMBER, they must take care to split the binning by sample. Any sequences from a different sample are missing from the reference and will be ignored, leading to incorrectly computed recall and precision.

To quantify the error, we compared recall and precision reported by AMBER for cross-sample binnings using single-sample references to the values when using a cross-sample reference with all sequences present. For 58 of the 203 bins ⋝ 200 kbp, the recall error was ⋝ 0.1, and for 39, it was ⋝ 0.5. For precision, the values were 86 of 203 and 34 of 203, respectively (Figure 4).

**Figure 4:**
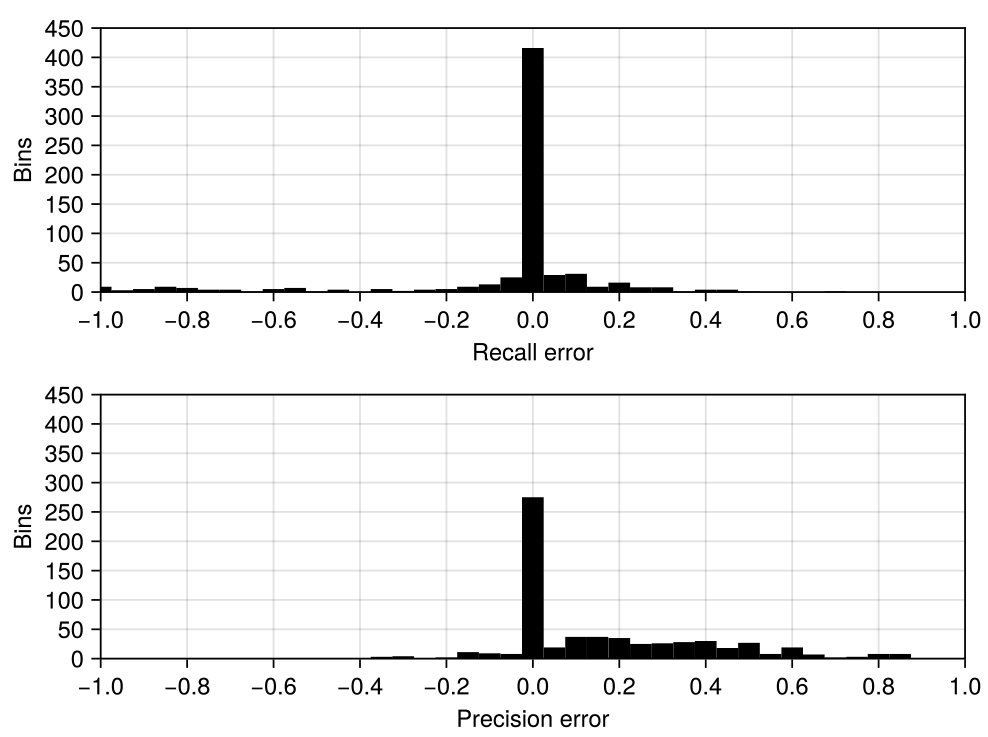
Benchmarking a multi-sample binning with a single-sample reference causes miscomputed precision and recall in AMBER due to sequences missing from the reference. The error reported here is the reported value minus the value when all sequences are present in the reference, for bins ⋝ 200 kbp in size.

While using AMBER with cross-sample binnings and single-sample references is a user error, it is an easy mistake to make. BinBencher will reject any binnings containing sequences not present in the reference, making this error impossible.

### 2.3. BinBencher accurately computes recall for poorly assembled genomes

When computing the recall of a genome/bin pair, it could be computed relative to the genome size (‘genomic recall’), or relative to the portion of the genome covered by any sequence input in the binner, typically the assembled part of the genome (‘asm recall’). Genomic and asm recall differ for genomes that are not wholly assembled and each may be the most useful metric depending on the objective. Let us define “recall gap” to be asm recall minus genomic recall.

The GI dataset is assembled with the CAMI ‘gold standard assembler’, a perfect ground-truth guided assembler, producing far higher quality contigs than what is realistically achievable using its input data[28], hence the recall gap for this dataset is small (Figure 5, left). To investigate the recall gap for more realistic data, we created a new ground truth reference by assembling the GI reads with the metaSPAdes assembler and binning the results (see Methods). For this dataset, 169 bins have a recall gap of ⋝ 0.1, and of the 49 bins with an asm recall of ⋝ 0.9, 22 had a genomic recall of < 0.5 (Figure 5, right).

**Figure 5:**
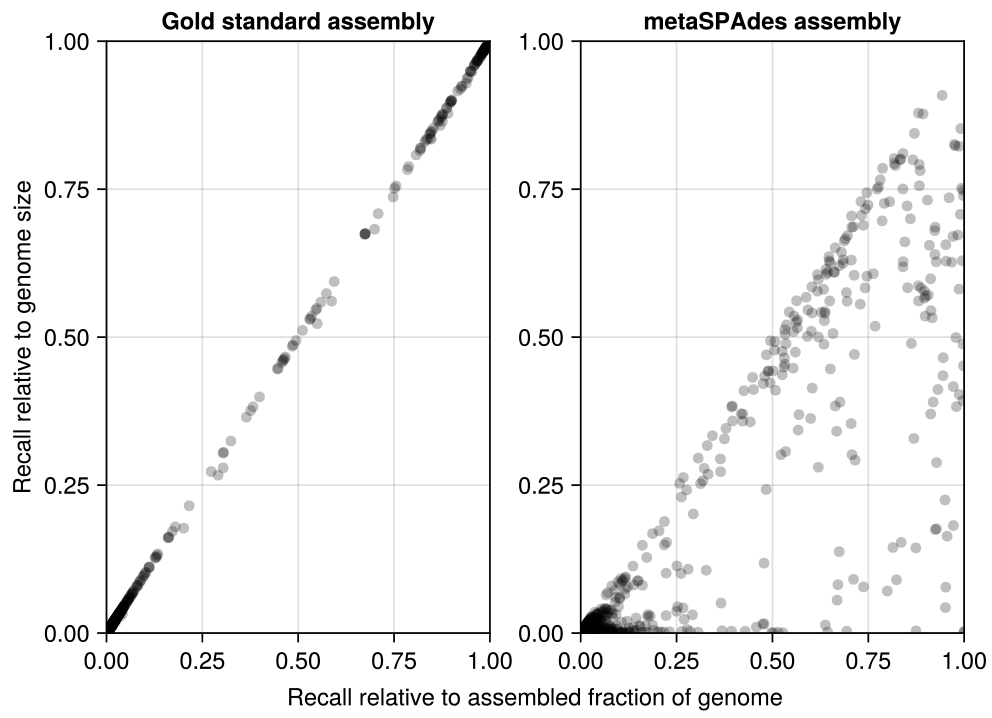
Recall relative to the assembled part of the genome (‘asm recall’), versus recall relative to the full genome (‘genomic recall’). Left: The GI gold standard assembly shows little difference between the two recall measures due to its unrealistically high quality. Right: A more realistic assembly using metaSPAdes reveals how genomic and asm recall may differ significantly for some bin/genome pairs.

Targeting asm recall when evaluating binning risks incentivizing underclustering, since small clusters are less likely to be contaminated, while still being able to encompass the assembled parts of genomes where only a few short sequences could be assembled. For example, in our experiment, bin 3993 had an asm recall of 1, because it consisted of the sole 4 kbp contig that was assembled from the bacterium *Taylorella equigenitalis*. Hence, the precision/recall tradeoff of a binner may be evaluated differently when assessing the number of high quality bins, depending on whether genomic or asm recall is used.

In our view, genomic recall aligns more closely to the common research objective of reconstructing whole and pure genomes. With this objective, asm recall can be misleading - a researcher might find a bin is assigned to some genome with a high recall, not realizing that this does not necessarily means the genome is recovered. On the other hand, since no binner can produce bins containing sequences that were not input to the program, asm recall has the useful property that the maximally achievable recall by any bin is 1, independent of how well the genome is assembled.

Because only reporting asm recall for poorly assembled bins can be misleading, BinBencher computes all statistics from both asm and genomic recall, but reports only those derived from the more biologically relevant genomic recall, unless explicitly requested.

### 2.4. Runtime and memory usage

To measure runtime and memory usage, we timed benchmarking all samples in the multi-split workflow. BinBencher ran faster than AMBER, taking 3 versus 180 CPU seconds, respectively. However, AMBER consumed only 267 MB memory compared to the 688 used to BinBencher.

**Table 1:**
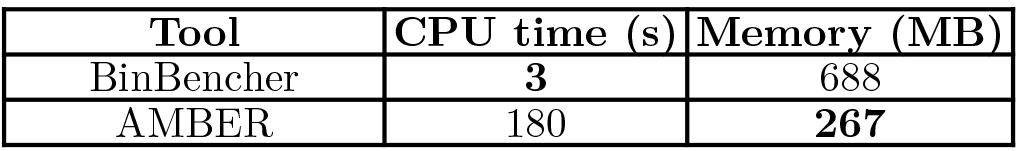
CPU and memory usage of binning benchmark tools.

## 3. Discussion

Evaluation is a necessary part of tool development. The chosen method of evaluation ultimately decides if a new tool, or a new development to an existing tool is considered an improvement. As we have shown in this paper, evaluating metagenomic binnings is not trivial, even with a ground-truth based reference available, but presents multiple pitfalls that can cause wrong or misleading evaluations. We believe these misleading evaluations can derail tool development. Indeed, during the development of our own binner, VAMB, we chased several promising techniques that turned out to only appear promising because of benchmarking artifacts. This experience made us invest significant amount of time in developing correct benchmarking techniques, eventually leading to the creation of BinBencher. We hope that the publication of BinBencher will enable other developers of binners to spend less time worrying about benchmarking, while also providing them with a more accurate metric to target.

Unfortunately, binning evaluation remains somewhat subjective. For example, there is no objective answer to how phylogenically precise a bin must be to be considered pure, nor an objective tradeoff between precision and recall. BinBencher strives to provide defaults that are generally biologically meaningful, and compute multiple metrics where there is no objectively right answer.

Nonetheless, BinBencher is still lacking in some important respects:

- BinBencher’s default measure, the number of recovered high-quality genomes, does not penalize the existence of additional poor quality bins. However, Bin-Bencher provide several additional metrics.
- BinBencher provides no option for handling chimeric contigs that ought to be split apart in multiple bins. This is because every sequence input to BinBencher must be present in the reference. However, we note that none of the binners we know about are able to detect and split chimeric contigs.

## 4. Methods

### 4.1. Computation of the default metric reported by BinBencher

BinBencher computes a variety of statistics, but the default reported metric is *number of recovered high-quality genomes*, which we define below.

The ground truth contains a number of genomes *G* that can be considered disjoint sets of genomic positions. *Y* is the set of all mapping positions, i.e. *Y* = ∪_*G*_. Let *X* be a set of sequences *S* to be binned. Each sequence *S* has length *L*_*S*_, and can be considered as a set of mapping positions *S* ⊆ *Y*, and its cardinality |*S*| may be larger, equal to, or smaller than *L*_*S*_. If we have a set of sequences *x* ⊆ *X* and a set of mapping positions *y* ⊆ *Y*, let us define *x* ⋒ *y* ≔ {*S* ∈ *x* | *S* ∩ *y* ≠ ∅}, the subset of *x* with sequences mapping to *y*.

A bin *B* is a set of sequences *B* ⊆ *X*. For any bin/ genome pair {*B, G*}, we have:

- TP_{B,G}_ = | ∪_*S*∈*B*_ *S* ∩ *G*|, is the true positives, the number of positions in *G* that any sequence in *B* is mapped to.
- FP_{B,G}_ = ∑_*S*∈*B*∖(*B*⋒*G*)_ *L*_*S*_ is the false positives, the sum of lengths of sequences in *B* that does not map to *G*.
- FN_{*B,G*}_ = |*G*| − TP_{*B,G*}_, the false negatives, are all positions in *G* not covered by any sequence in *B*.

From these definitions we define recall *R*_{*B,G*}_ and precision *P*_{*B,G*}_ the usual way. We can then count the number of recovered genomes at recall/precision thresholds *T*_*R*_, *T*_*P*_ as 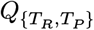 =

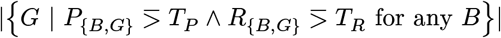

The default thresholds are *T*_*R*_ = 0.9, *T*_*P*_ = 0.95.

There are several differences between BinBencher’s and AMBER’s computed measures, but the most important is that AMBER uses TP_{*B,G*}_ = ∑_*S*∈*B*⋒*G*_ *L*_*S*_ and FN_{*B,G*}_ = ∑_(*X*⋒*G*)∖*B*_ *L*_*S*_, i.e. it counts the sum of the lengths of sequences instead of unique mapping positions.

### 4.2. Benchmarking on multiple phylogenetic ranks

In BinBencher reference files, every genome is organized in a phylogenetic tree. Genomes are the lowest taxonomic rank in the tree, assigned rank zero. Every member of taxonomic rank *T* has a parent of rank *T* + 1, except the final taxonomic rank which has no parent and only one member, the ancestor of every genome in the reference.

BinBencher computes precision/recall for every {*B, C*_*T*_ } pair of bin with a clade of rank *T*. The values of {*B, C*_0_} are bin-genome pairs, and their precision and recall are computed as shown in the previous section.

We denote the set of direct children of clade *C*_*T*_ to be *H*(*C*_*T*_). For the non-genomes clades *C*_*T*_ where *T* > 0, we have 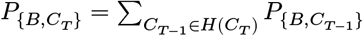 and *R*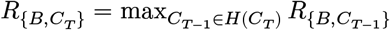.

### 4.3. Dataset and software used

For the results presented in this paper, we used the 10 synthetic Gastrointestinal short-read samples from the 2nd CAMI Toy Human Microbiome Project Dataset (the “GI” dataset)[29]. We ran VAMB v4.1.3 to produce the bins measured in this manuscript, and benchmarked the output using BinBencher v0.3.0 and AMBER v2.0.4

### 4.4. Reassembling and binning the GI dataset

To assemble the GI dataset, we ran SPAdes v. 3.15.5 with the – `meta` option[33] and default parameters on reads from each sample. We then concatenated output contigs of length ⋝ 2 kbp to a single file used as input to binning. To get a reference, we aligned contigs from each sample against the underlying genomes which had a nonzero abundance in that sample using BLAST v2.15.0 [35], and accepted all hits with the arbitrary cutoffs ⋝ 95% identity and ⋝ 90% query coverage. The re-assembled contigs were binned with VAMB v4.1.3 using default parameters.

## 4.5. Code availability

The code used to produce this paper can be found at https://github.com/jakobnissen/paper_binbencher. BinBencher itself is freely available at https://github.com/jakobnissen/BinBencher.jl.

## 4.6. Data availability

The GI dataset is made available by the Critical Assessment for Metagenome Evaluation, at https://frl.publisso.de/data/frl:6425518.

## 4.7. Acknowledgements

This work is supported by the Novo Nordisk Foundation (NNF20OC0062223 and NNF23SA0084103).

